# Spatial organization of mediated-macrophage chemoprotective niches in solid tumors: A mathematical analysis

**DOI:** 10.1101/2024.11.21.624654

**Authors:** William Dangelser, Angélique Stéphanou, Arnaud Millet

## Abstract

Acquired resistance is one of the major causes of failure of standard therapies in cancer patients. Chemotherapeutic agents are still widely used and the understanding of the mechanisms leading to secondary resistance to these molecules are still puzzling. Recently, the role of the tumor immune microenvironment has been recognized. Among the cells potentially involved, macrophages seem to be the perfect culprits. In a previous work, we have shown that hypoxic macrophages are able to provide strong protection against 5-fluorouracil, a first-line chemotherapeutic agent in digestive cancers. In the present work, we use mathematical modeling to explore the spatiotemporal aspects of the treatment-induced organization of the tumor environment. Based on analytical and numerical analysis, we propose that macrophage-driven protection against chemotherapy under treatment does not rely solely on biochemical degradation, but is enhanced by the emergence of spatially structured chemotherapeutic protective niches. This work paves the way for the development of new therapeutic strategies that rely on targeting the spatial organization of tumors as a way to control treatment resistance.

## 1 Introduction

Solid tumors are a leading cause of death worldwide. Despite recent advances in the use of innovative therapeutic strategies, chemotherapy remains the current standard of care for these diseases. Chemotherapies have been developed for more than half a century and are based on their ability to interfere with the proliferation and survival of cancer cells. Chemotherapeutic molecules are classified according to their mechanism of action: alkylating agents, antimetabolites, anti-microtubule molecules, ribonucleic synthesis inhibitors and others such as proteasome inhibitors (Goodman et al (1996)). Since their discovery, the use of these molecules has been confronted with the emergence of resistance. Resistance to chemotherapy involves several mechanisms. Some may be related to the molecule itself, such as a decrease of drug uptake, an increase in drug efflux, sequestration of the drug or an alteration in drug metabolism. Others are related to the target cell such as an increased capacity for DNA repair, an inhibition of apoptosis induction and the induction of a protective autophagy. In addition, the physics of the environment may be involved, such as the spatial organization of the vascular network or the composition of the extracellular matrix, which may induce changes in the hydrodynamic resistance of the tissue and thus interfere with the penetration of the drug into the tissue (Tyner et al (2022)).

In recent years, the immune system has been recognized as a key element in understanding the mechanisms of tumor interaction with surrounding healthy tissues, as well as a provider of new therapeutic strategies. The tumor immune microenvironment (TIME) is composed of different types of immune cells. However, tumor associated macrophages (TAMs) typically represent the quantitatively largest population found in solid tumors. TAMs are involved in tumor growth, immune evasion, neo-angiogenesis and treatment resistance. Using depletion methods, a large number of studies have reported an increased chemosensitivity when macrophages are removed from the tumor (Ruffell and Coussens (2015)). Furthermore, co-culture studies have revealed macrophage-mediated resistance mechanisms to various anticancer drugs such as paclitaxel, doxorubicin, etoposide or gemcitabine (Mitchem et al (2013), Shree et al (2011)). Specifically, depletion of MHCII^*lo*^ TAMs results in increased sensitivity to taxol-induced DNA damage and apoptosis (Olson et al (2017)). Mechanisms involving macrophage-induced chemoresistance typically rely on macrophage secretion of factors such as pyrimidine nucleosides (deoxycytidine), which inhibit gemcitabine induction of apoptosis in pancreatic ductal adenocarcinoma (Halbrook et al (2019)). In colorectal cancer, for example, the involvement of macrophages in chemoresistance has been suggested based on *in vitro* and *in vivo* studies. The proposed mechanisms are diverse but mainly involve secreted factors and are quantitatively moderate. For example, it has been proposed that IL-6 secreted by macrophages can stimulate STAT3 in cancer cells, which induces the inhibition of the RAB22A/BCL2 pathway through miR-204-5p expression, thereby leading to chemoresistance to 5-fluorouracil (5-FU) (Yin et al (2017)). Similarly, macrophage secretion of putrescin, a member of the polyamine family, has been shown to suppress the JNK/Caspase 3 pathway in cancer cells, conferring protection against 5-FU (Zhang et al (2016)).

Recently, we proposed that macrophages may massively contribute to chemoresistance to 5-FU, a pyrimidine analog drug with antimetabolite activity, in the context of colorectal cancer. Indeed, we have shown that human macrophages in low oxygen environments display a strong expression of the dihydropyrimidine dehydrogenase (DPD), an enzyme responsible for the first step of the pyrimidine bases catabolism pathway, capable of metabolizing 5-FU to an inactive product. In this work, we have shown that this mechanism is specific to human macrophages and favors a global tumor resistance without the involvement of cancer cells resistance (Malier et al (2021)). The specific expression of DPD in tumor associated macrophages has been confirmed in an independent study (Cui et al (2023)) and the clinical relevance of this finding is currently under investigation. Despite these results, the prediction of treatment resistance in a peculiar patient is still difficult. One reason for this is that we lack a quantitative understanding of the dynamics of the interaction between cancer cells and macrophages in the context of a chemotherapy regimen. To advance this understanding, we developed mathematical models, both analytical and numerical, to study the spatiotemporal dynamics of a tumor microenvironment subjected to a chemotherapy.

## 2 Continuous Model

In this section, we describe the continuous model used in this study. Macrophages, cancer cells, a chemoattractant secreted by cancer cells (e.g. CSF1) and a chemotherapy (e.g. 5-FU) are characterized by their spatial distribution which is assumed to be continuous and smooth. Cells and molecules are submitted to passive diffusion. Macrophages are sensitive to chemokine-mediated attraction released by cancer cells and cancer cells are sensitive to chemotherapy. To take into account the role of macrophages in chemoresistance, we introduce a macrophage-dependent degradation of chemotherapy as illustrated in Figure 1. Let consider the tissue space as a smooth bounded connected domain of ^2^ denoted Ω. Macrophage and cancer cells densities are represented by *m*(**x**, *t*) and *k*(**x**, *t*) respectively and similarly concentrations of chemotherapy and chemoattractant are denoted by *c*(**x**, *t*) and *a*(**x**, *t*). Both cancer cells and macrophages present a random migration that will be associated with a constant diffusion coefficient *d*. Furthermore, macrophages present also a directed migration behavior directed by the gradient of secreted chemoattractant. This chemoattraction is introduced in the model by a chemotaxis coefficient χ that could depend on the macrophages density. Cancer cells are supposed to follow a logistic growth pattern with a growing rate *α* and the carrying capacity denoted *K*. Chemoattractant and chemotherapy are considered free molecules in Ω and then subjected to Brownian diffusion characterized by a constant diffusion coefficient *D*. Chemotherapy interferes with cancer cell growth with a chemotherapy-induced death rate *β*. The *γ* term represents the experimental finding that DPD expression in macrophages leads to 5-FU conversion to 5-FUH_2_, an inactive product. Chemotherapy is introduced in the model by a source term *s*(**x**, *t*). Finally, chemoattractant is secreted by cancer cells at a rate *δ* and present a spontaneous degradation rate *η*.

**Figure 1:**
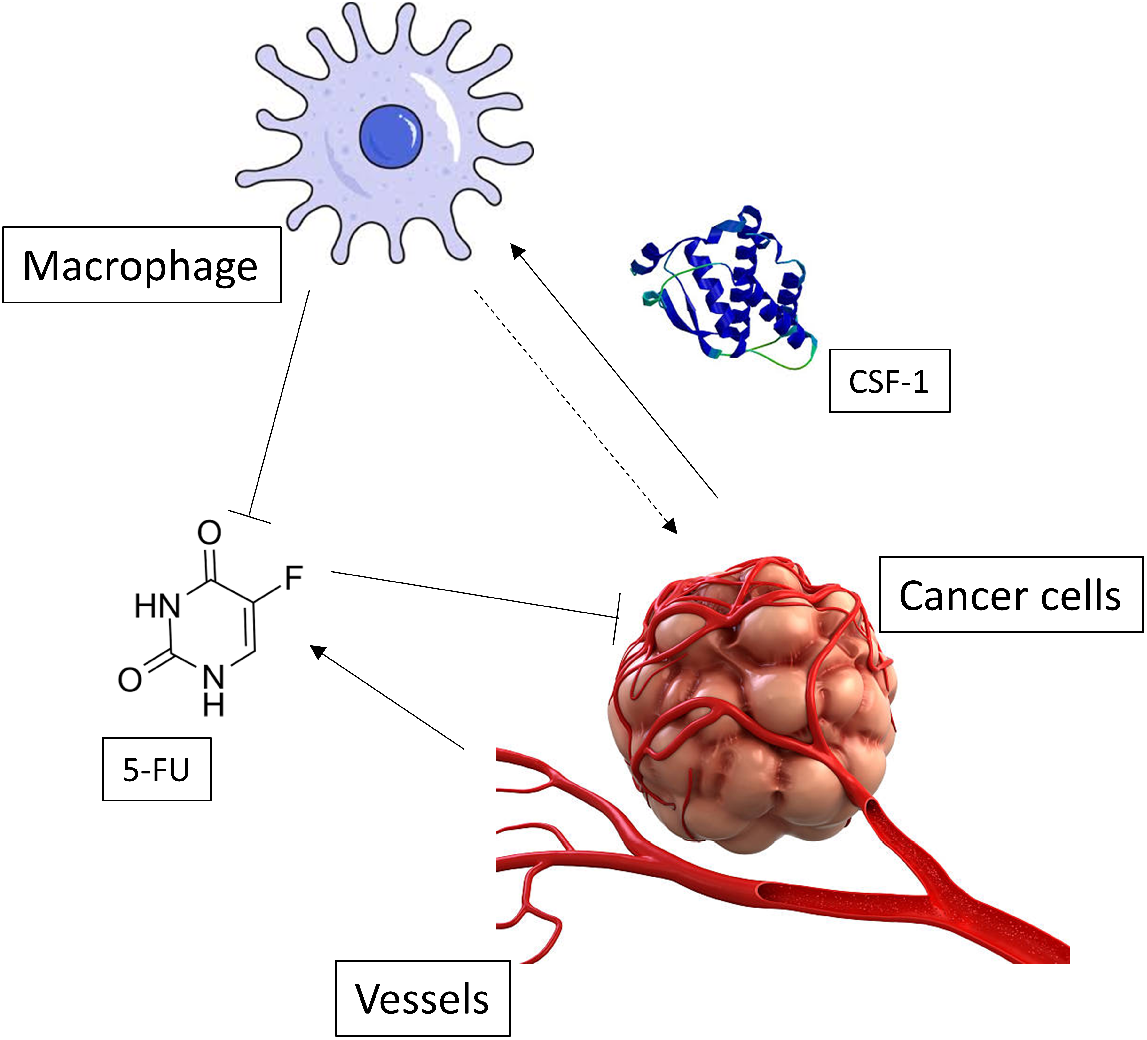
Schematic of the model

The equation system associated with this model reads as follows:

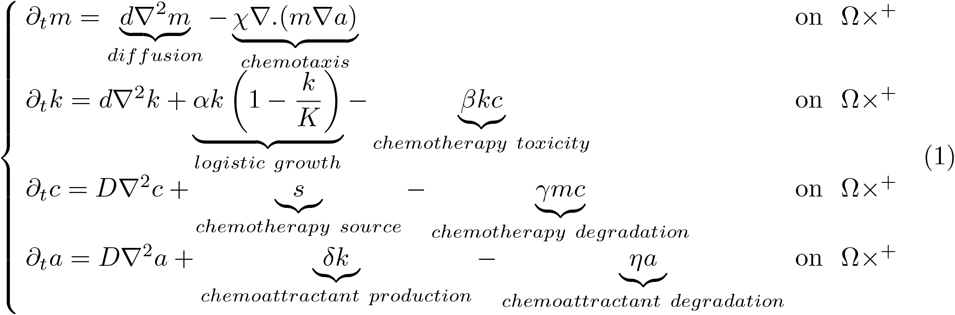

### 2.1 Nondimensionalization

In order to define dimensionless variables we introduce the following rescalings :

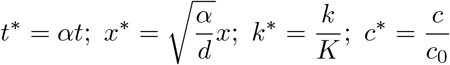

Choosing *K* as a density reference for cells and *c*_0_ for chemotherapy and chemoattractant, the other variables and parameters of our system are:

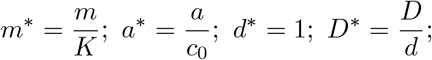

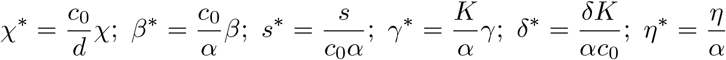

Dropping the asterisks, the system (1) takes the following nondimensional form

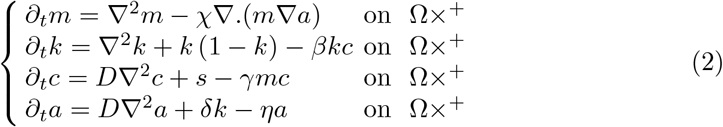

### 2.2 Spatial patterning conditions

#### 2.2.1 Homogeneous steady states

First, we consider the system (2) omitting the spatial dependence in order to study the spatially homogeneous steady states and we consider the source of chemotherapy as a constant in time and space. We find two steady states: 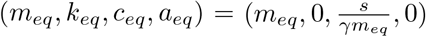 or 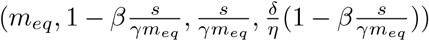. The first solution is cancer free so we will consider only the second solution in the following. This second solution exists only if 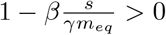. Introducing

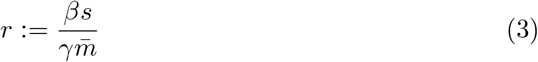

we can express our condition under the simplified following form

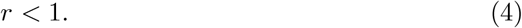

A linear analysis of (4) demonstrates that the steady state 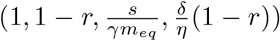 is locally asymptotically stable.

#### 2.2.2 Linear analysis

We note 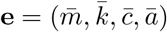 the stable steady state for which the previous assumption (4) is fulfilled. In order to study the spatial stability, we linearize (2) around **e**. Writing *W* := (*m, k, c, a*) the perturbation of **e**, we obtain the following linearized system

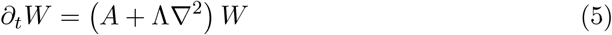

where

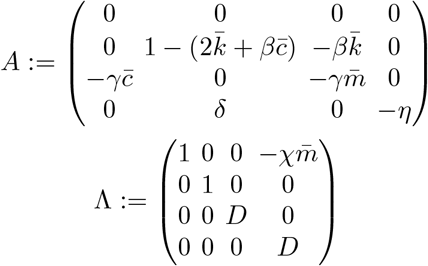

Looking for solutions of the following form **u***e*^*λt*+*iκ*.**x**^ where *κ* refers to the wave vector. Noting

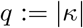

We get from (5)

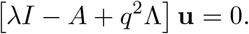

The determinant of this matrix will give us conditions for the existence of spatial instabilities. Indeed, the determinant could be expressed as:

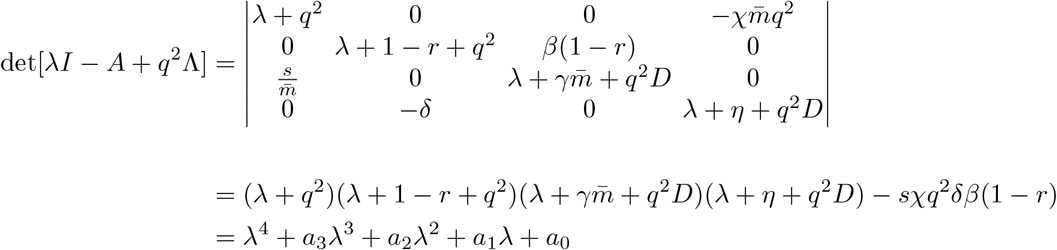

where the a factors are detailed in the appendix A.

According to the Routh-Hurwitz criterion, a sufficient condition for instabilities is *a*_0_(*q*^2^) < 0 for some *q*, so:

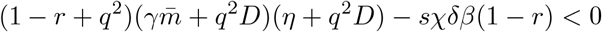

We develop this expression as a polynomial of degree three in the variable *q*^2^:

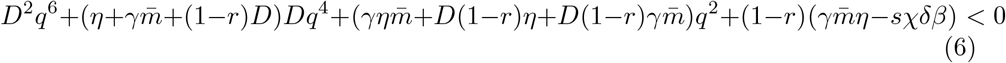

As we know that every coefficients preceding power terms of *q*^2^ are positive, the Routh-Hurwitz condition (6) is equivalent to

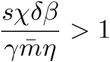

Finally our stability analysis provides the following condition for the existence of spatial patterning constraining the regime of chemotherapy source as

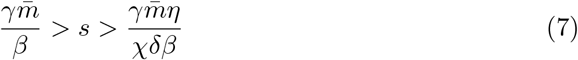

This result illustrates the existence of a spatial organization driven by the introduction of chemotherapy but in a specific range of amplitude. Too few chemotherapy is unable to drive the organization as macrophages degrade it to rapidly and too much the mechanism of degradation is overwhelmed by the chemotherapy. Using physiological order of magnitudes, we can estimate the values of the range of chemotherapy necessary to drive spatial organization. Indeed, using the values of parameters reported in Appendix B we can estimate the upper bound of the interval of *s* to be 10^12^ and its lower bound to be 10^1^. The amount of 5-FU measured in tissues from patients suffering a colorectal cancer corresponds to a value *s* ≈ 1 10^−2^ g*/*m^3^ s which in its nondimensionalized form is 10^10^ fulfilling the conditions (7). This results suggests that a spatial organization induced by 5-FU is physically possible in real tumors.

### 2.3 Numerical analysis

In that section, we describe the numerical analysis of the continuous model described in previous sections. The following simulation results were obtained by solving the system (2) with an explicit scheme.

#### 2.3.1 Initial conditions and boundary conditions

To simulate the spatiotemporal dynamic of instabilities, we start from the equilibrium found in section 2.2.1 and we add a perturbation using a white noise over imposed on cancer cells. The corresponding initial conditions are

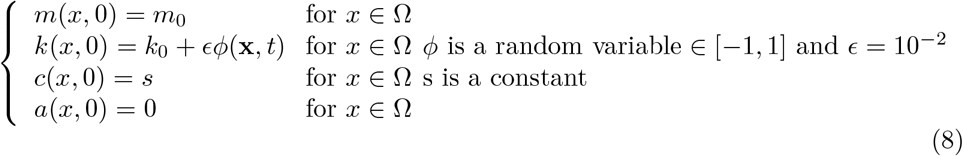

We impose reflecting boundaries at the frontier of our domain Ω which corresponds to Neumann conditions:

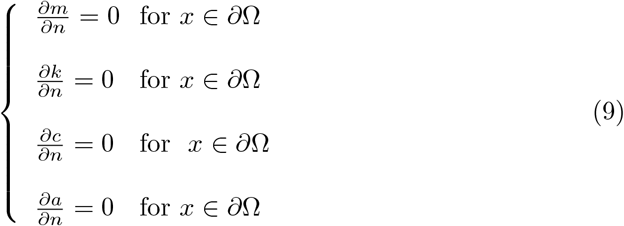

#### 2.3.2 Spatial organization of macrophages driven by chemotherapy

Numerical simulation reveals that when the chemotherapy source term *s* fulfills the condition (7), patterns appear after a small perturbation of the cancer cells distribution after 20 days (Figure 2). Of note, the macrophage and cancer cell patterning is driven by the chemotherapy as the *s* = 0 condition does not lead to any patterns (Figure (3), as expected according to (7)). The qualitative behavior of patterning is insensitive to the level of chemotherapy as long as condition (7) holds, as it is illustrated in Figure 4 where increasing values of *s*, from the same initial conditions, only accelerates the formation of patterns without modifying their shape. To assess the stability of the patterns, we have simulated the effect of chemotherapy for 10 days (Figure 5a) and then simulate the dynamic when chemotherapy is maintained (Figure 5b) or stopped (Figure 5c) until 20 days. This dynamical analysis shows that macrophage patterning induced by the first round of treatment is still present after a similar period of time than the exposure to chemotherapy. This result illustrates the possibility to have a tissue memory of a previous treatment round through the spatial organization of macrophages. This suggests that a second round of treatment will face a spatially organized tissue more prone to resist to its action than it was initially the case. To further explore the impact of the formation of macrophage-mediated chemoprotective niches we will use an agent-based model in the next section.

**Figure 2:**
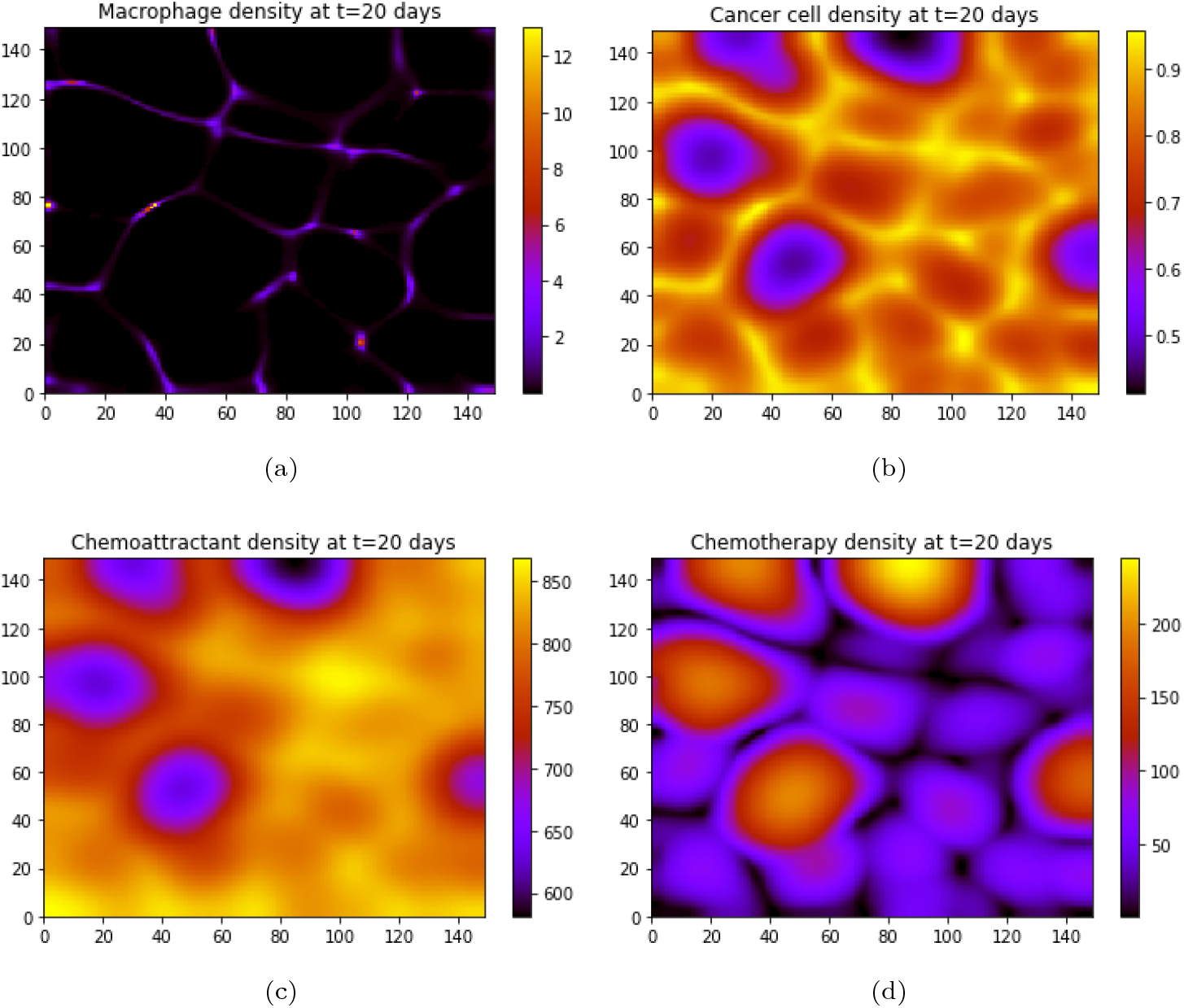
Simulations of the continuous model after 20 days. The parameters used are *D* = 10^4^, χ = 10, *m*_0_ = 0.1, *β* = 10^*−*2^, *s* = 10^4^, *γ* = 10^4^, *δ* = 10^5^, *η* = 10^2^. The value of *s* respects the instability condition (7).

**Figure 3:**
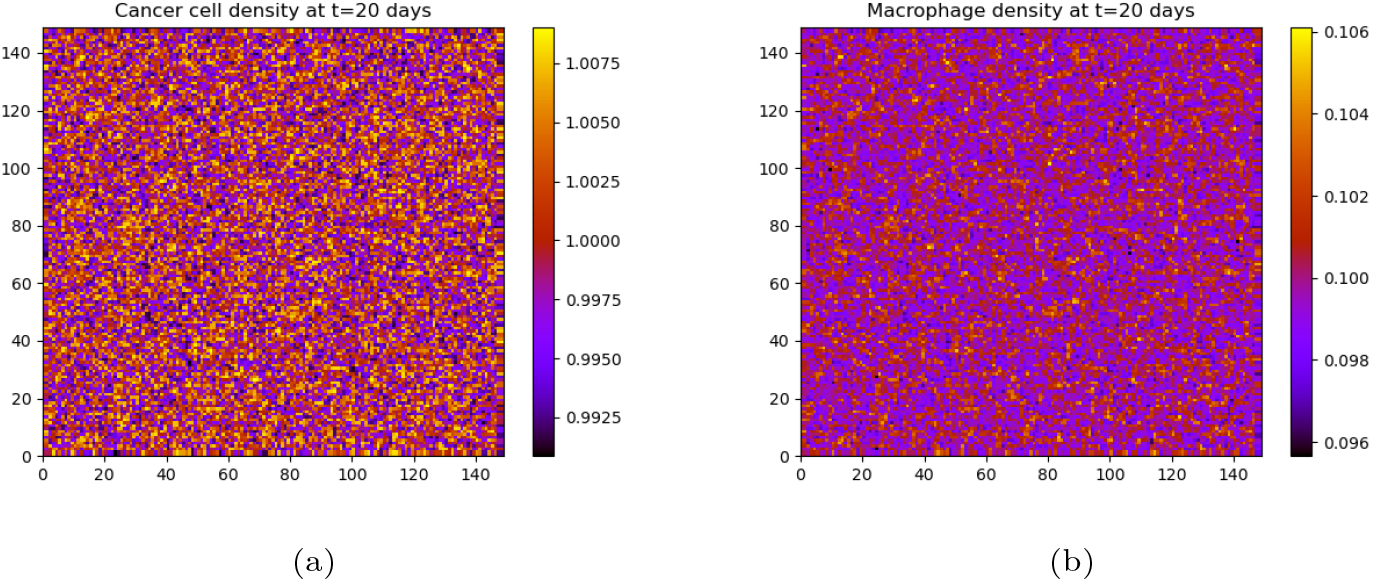
Simulations of the continuous model without chemotherapy after 20 days. The others parameters are the same as in Figure 2. (a) cancer cells, (b) macrophages.

**Figure 4:**
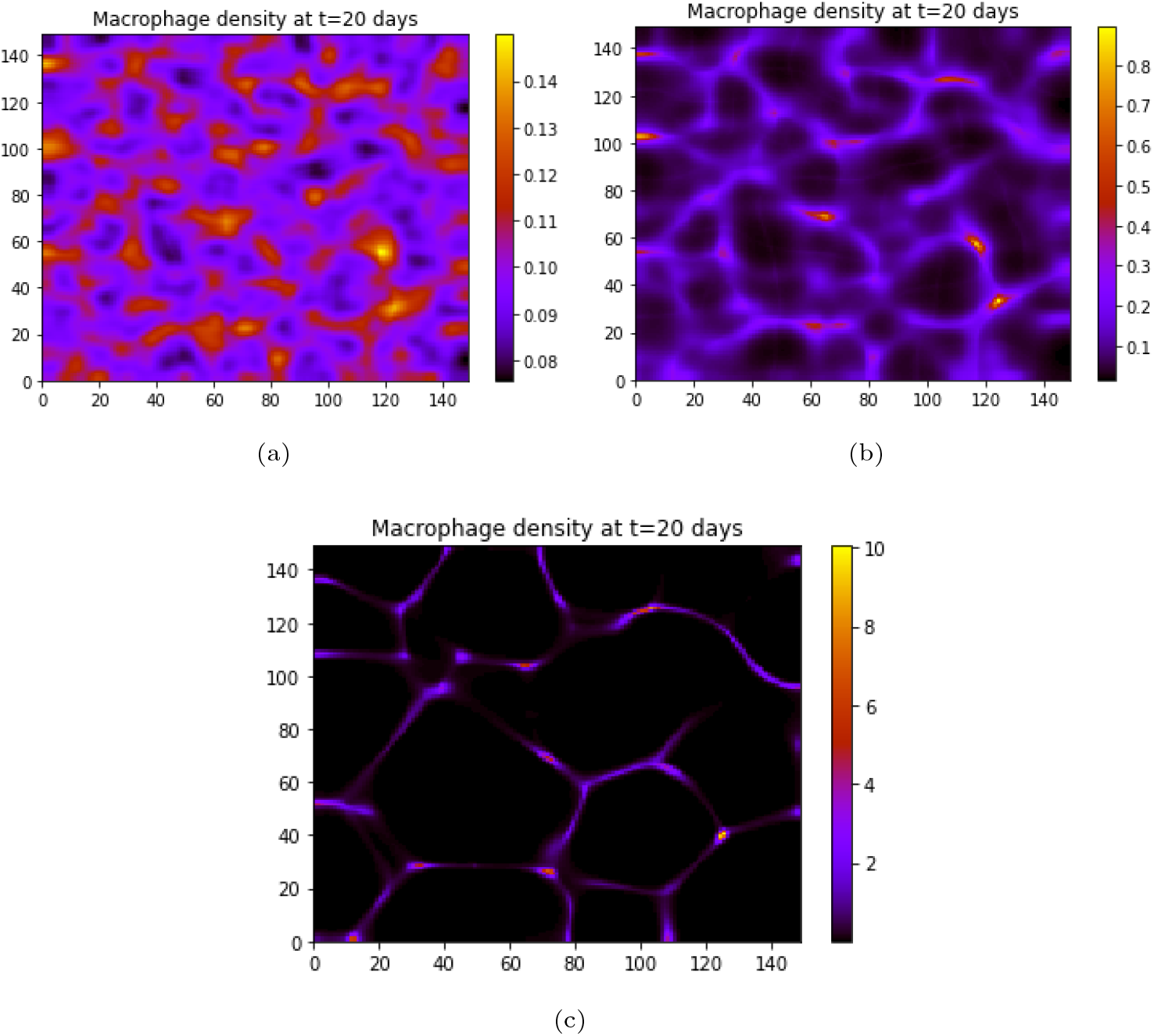
Simulations of the continuous model for different values of *s* after 20 days. The others parameters are the same as in Figure 2. (a) *s* = 10^3^, (b) *s* = 5 × 10^3^, (c) *s* = 10^4^. These values still satisfy condition (7).

**Figure 5:**
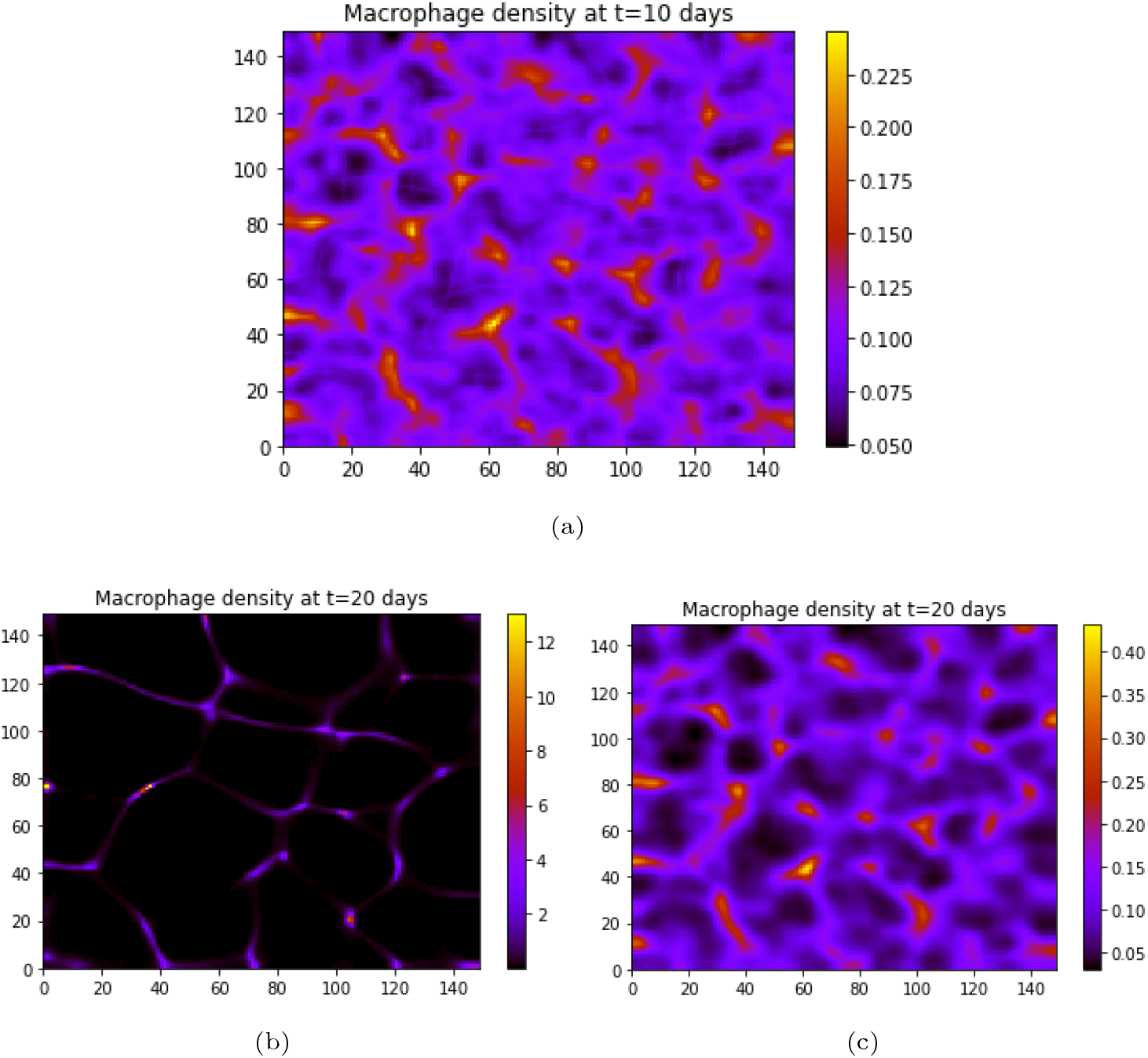
Simulations of the continuous model (a) after 10 days, (b) after 20 days without stopping the drug influx, (c) after 20 days and the drug influx was stopped after 10 days. The parameters are the same as in Figure 2.

**Figure 6:**
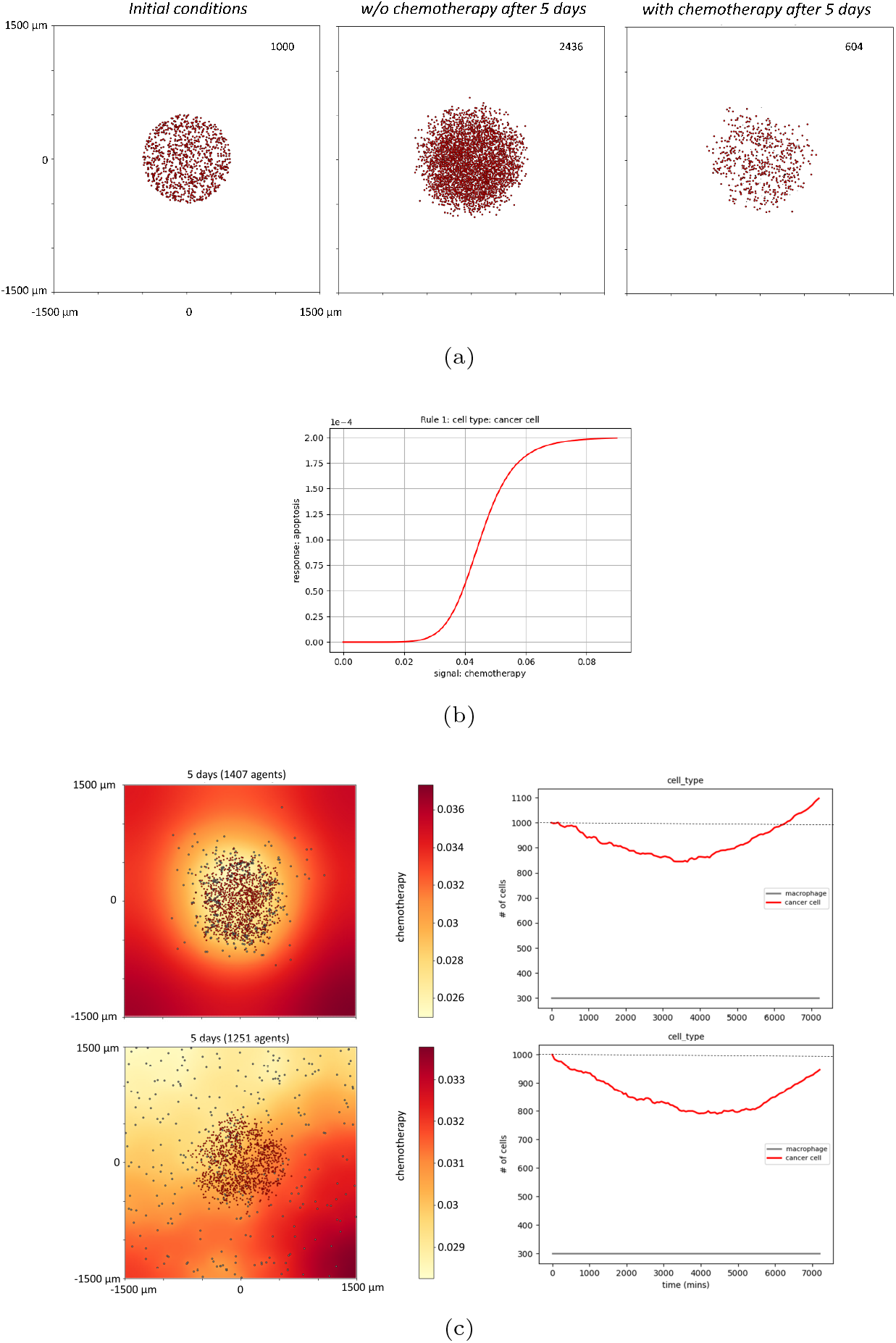
Agent-based simulations. (a) Left panel represents the initial conditions of cancer cells, the middle panel shows the growth and diffusion of the cancer cells without chemotherapy after 5 days and the right panel shows the same evolution but under treatment during the same period of time (the final number of cancer cells is indicated in the upper right corner of each panel). (b) Representation of the Hill curve corresponding to the cancer cell response to chemotherapy. (c) Evolution of the population of cancer cells and macrophages after 5 days when macrophages are attracted by cancer cells (upper panel) and when no chemoattractant is secreted by cancer cells (lower panel). The right panels shows the temporal evolution of the two populations compared to the initial conditions (dashed line).

## 3 Agent-based simulation

The continuous model used in previous sections relies on the hypothesis that our analysis takes place at a spatial precision where cell size is negligible. This approach presents the advantage to authorize an analytical resolution and to give us an intuition of what happens in the tumor microenvironment. Nevertheless, the spatial organization of tumors could present functional structures involving few cells as it is the case in precancerous states or metastasis where this continuity hypothesis no longer stands. There is a clear advantage in this case to propose an analysis where each cell has its own identity and to abandon the cell density point of view. In order to perform this task, we will use an agent-based model. We chose the physics-based multicellular simulator *PhysiCell* (Ghaffarizadeh et al (2018)) and more specifically the *PhysiCell Studio* version that provides a graphical user interface (Heiland et al (2023)). *PhysiCell* is an open-source agent-based simulator coded in C++ designed for multicellular systems. It authorizes to create two kinds of objects: substrates and agents. Substrates are computed on a Cartesian mesh and are mostly used to model all the chemicals in the cellular microenvironment, in other words the continuous components of the model. In our case, we will use this approach for the chemotherapy and the chemoattractant. Agents consist of spheres (3D) or disks (2D) that are described by their center and their volume on a specific grid. Multiple types of cells can be computed in the same model and each one has its own properties. *PhysiCell* allows the user to set the volume, death, type of cycle for proliferation or motility parameters of the cells and their interactions with the substrates. Using a PDE-solver, the following equation for the substrates is solved at each time step

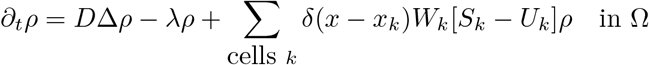

where *ρ* is the vector of substrates, *D* and *λ* are the vectors of diffusion coefficients and decay rates respectively, *δ* is the Dirac delta function, *x*_*k*_ is the *k*^*th*^ cell position, *W*_*k*_ its volume, *S*_*k*_ and *U*_*k*_ are the source and uptake rates.

### 3.1 Agent-based simulation of chemo-protective action of macrophages

Starting from a random distribution of 1000 cancer cells located in a circle of radius 500 *µ*m embedded in a computational domain Ω of 1500 × 1500 *µm*^2^ with Neumann conditions applied at its boundary *∂*Ω, we introduce a chemo-attractant secreted by cancer cells. The chemotherapy is initially homogeneously distributed in the domain Ω. Cancer cell growth is parameterized with the same *α* used in the continuous model (Appendix B) (Figure 6a). Cancer cell response to chemotherapy is modeled by a Hill curve (Figure 6b) resulting in apoptosis (as shown in the right panel of Figure 6a). We then introduce a spatially randomly distributed population of macrophages maintained constant in number during the time experiment (N=300), according to the suspected low division rate of human macrophages *in vivo*. The rate of chemotherapy degradation by macrophages was determined thanks to a simulation of macrophages alone. The numerical values were set to obtain the same half-life as that observed experimentally (Malier et al (2021)). This approach is based on the hypothesis that chemotherapy degradation follows a first-order decay law. This hypothesis is justified when macrophages presents a low density in the continuous model as shown in Appendix C.

The simulation shows the ability of macrophages submitted to chemo-attraction by cancer cells to protect tumor cells from chemotherapy resulting in an increase of the population number of cancer cells after 5 days (Figure 6c). Interestingly, without chemoattraction the same population of macrophages is not associated with the growth of the tumor at 5 days. This result reinforces the conclusion obtained by the continuous model revealing the importance of the chemoattraction of macrophages to provide an efficient protection along with a spatial organization preventing future efficacy of the treatment.

### 3.2 Spatial organization of macrophages determines the fate of the tumor after multiple cycles of treatment

To further evaluate the importance of the spatial organization of the macrophages/-cancer cells system, we simulate a treatment protocol comprising three cycles of 5 days of chemotherapy diffusing from a vessel. The diffusion of the chemotherapy from a vessel authorizes to study the impact of a symmetry breaking in the model (Figure 7a) contrary to the homogeneous initial condition used in Figure 6. In this model, macrophages confer a resistance toward a treatment regimen able to block tumor growth effectively (Figure 7b and 7c). Preventing the chemoattraction of macrophages by cancer cells and maintaining their ability to metabolize the chemotherapy leads to a less effective protection (Figure 7d). This result illustrates the importance of the spatial organization of the innate immune response in resistance to chemotherapy.

**Figure 7:**
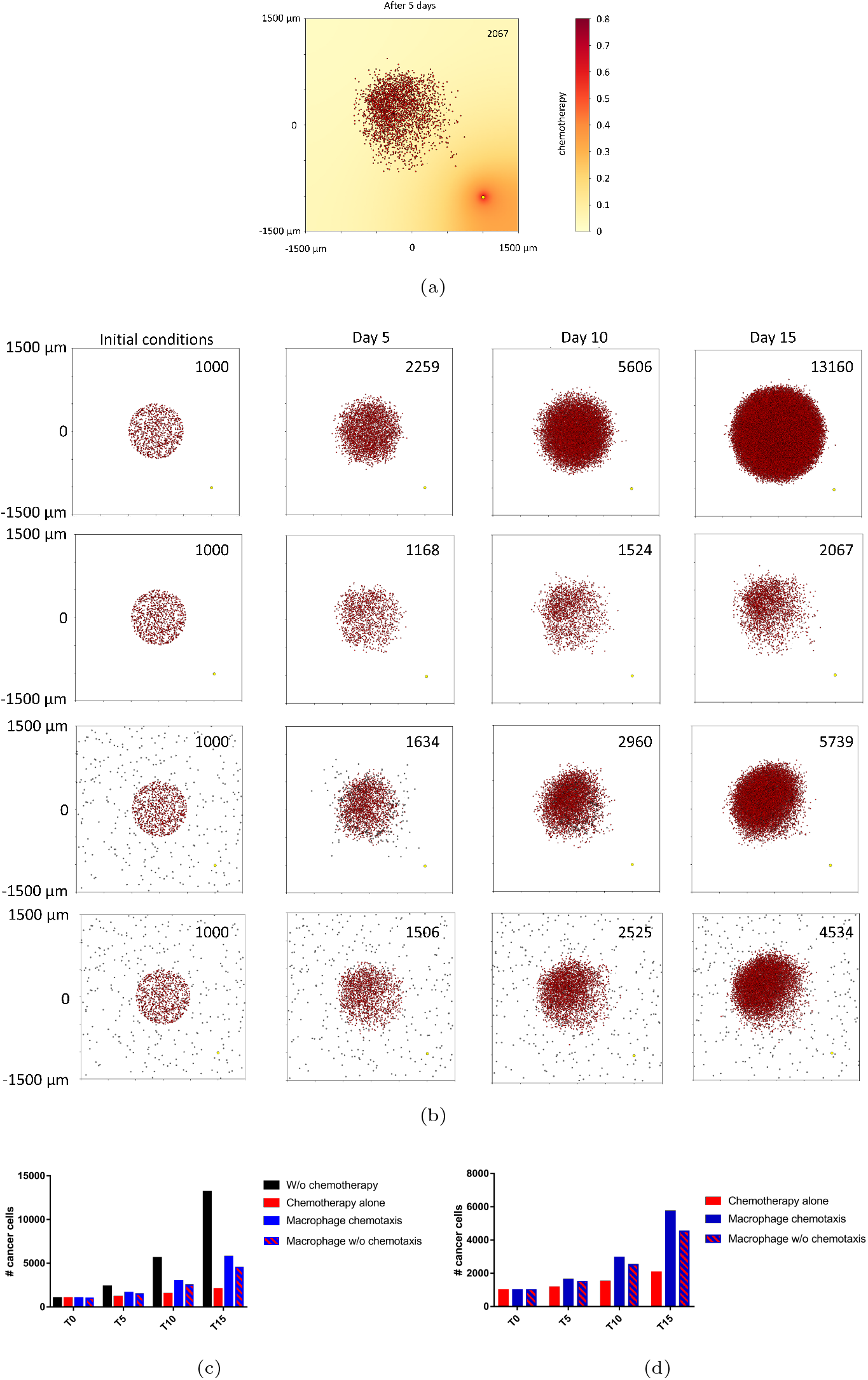
Simulations (a) representing the evolution of a tumor after 3 cycles of chemotherapy diffusing from a vessel located at the bottom right of our space, (b) representation of the evolution during the three cycles of the tumor w/o chemotherapy, with chemotherapy, with macrophages and with macrophages w/o chemoattractant from top to bottom respectively (the final number of cancer cells is indicated in the upper right corner of each panel), (c) evolution of the population of cancer cells in the four conditions at the end of each treatment cycles, (d) evolution of the population of cancer cells only for conditions associated with treatment.

## 4 Discussion

The spatial heterogeneity of tumors has been recognized for a long time, but due to the lack of adequate technological tools, the specific study of the 3D organization of tumors has not received the attention it deserves. This situation is changing, especially with the use of new spatially resolved techniques to study the expression of proteins, mRNA or metabolites. More importantly, the spatial organization of the tumor-associated immune response is of particular interest since the advent of the use of immunotherapies to predict their efficacy (e.g. anti-PD-1 antibodies) (Di Mauro and Arbore (2024)). However, the direct involvement of immune cells in tissue organization has received little attention, even though macrophages represent the largest population of immune cells in many solid tumors. These innate immune cells are present in every tissue as resident cells and could be recruited from circulating monocytes during inflammatory processes. Despite their potential to fight cancer cells, in many tumors macrophages are known to be part of the supportive environment that facilitates tumor growth. In this context, they are termed tumor-associated macrophages (TAM) to highlight their pro-tumor role. Recently, TAMs have been identified as a source of treatment resistance to chemotherapeutic agents, but also to immunotherapies (Ruffell and Coussens (2015), Bader et al (2024)). The reported mechanisms are usually analyzed at the molecular level and provide little information on a potential spatial component of the resistance. Experimentally little is known about the active participation of macrophages in the spatial organization of the tumor despite their ability to remodel the extracellular matrix (Bied et al (2023)). Some studies have explored this issue at the mathematical level, a seminal paper by Owen and Sherratt suggested that the secretion of a chemotactic molecule by cancer cells could lead to spatial irregularities in the distribution of macrophages (Owen and Sherratt (1997)). However, the importance of macrophages in the spatial organization of tissues under treatment has not been addressed. The demonstration that macrophages could metabolize by direct contact a chemotherapeutic agent (5-FU) resulting in a resistance of the tumor without any intrinsic resistance of cancer cells forced us to analyze the implication of the spatial organization of protective areas, called chemoprotective niches. To study the role of macrophages in this context, we analyzed the simplest model involving cancer cells, macrophages, a chemoattractant molecule (e.g. CSF1) and a chemotherapy (e.g. 5-FU) (Figure 1).

The analytical condition (7) and numerical analysis revealed the existence of a range of chemotherapy, corresponding to physiological relevant levels, leading to spatial heterogeneity forming protective niches. According to our model, two strategies could be proposed to deal with this spatially driven resistance. First, we can try to target the biochemical degradation process, which in our model means decreasing *γ* and/or 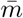. This can be done by using a specific inhibitor of DPD, such as gimeracil. This strategy has already been proposed using a combination drug containing 5-FU and gimeracil under the name S-1 (Sakuramoto et al (2007)). Despite promising preliminary results, this approach has not revolutionized the standard of care for patients with digestive cancers. The main reason is probably the uncontrolled distribution of gimeracil without specific targeting of the tumor. Another possibility is to reduce the number of macrophages by targeting the CSF1-R receptor with a specific antagonist (Cannarile et al (2017)). In fact, this strategy will not lead to a reduction of macrophages in humans, but to a reprogramming of macrophages leading to a reduction of DPD expression under the control of HIF-2*α*, as we have recently shown (Gharzeddine et al (2024)). The second strategy is to reduce the production rate *δ* of chemoattractant or macrophage sensitivity χ to that chemoattractant, in our case CSF1. Again, a CSF1-R antagonist seems to be the best candidate as it provides the expected double-effect to lower DPD activity and prevent chemoattraction toward cancer cells. An intriguing result of the present work is that this combined strategy will provide the best strategy to reduce the possibility of inhibiting the formation of chemoprotective niches. In this case, the upper bound of (7) will indeed be lowered. Indeed, if the effect of the drug is only to decrease *γ*, the lower bound will also decrease, leading to the possibility of patterning at lower 5-FU concentrations. Synchronous targeting of the χ term and the *γ* term, which is achieved by CSF1-R targeting, will induce a narrowing of the patterning conditions (7).

From this we can conclude that CSF1-R targeting is probably one of the best strategies to prevent any chemotherapy-induced spatially organized resistance in a macrophage-rich tumor microenvironment.

## Acknowledgements

AM is supported by the Ligue régionale contre le cancer and Cancéropole Auvergne Rhône Alpes CLARA.

## Declarations

- Data availability The research materials supporting this publication can be accessed by contacting

## Appendix A

The terms of the polynomial corresponding to the matrix of the linear analysis section 2.2.2 are

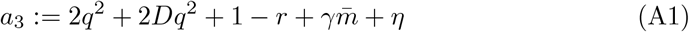

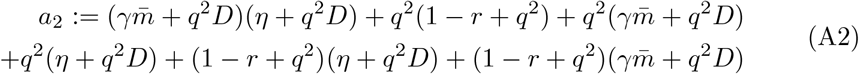

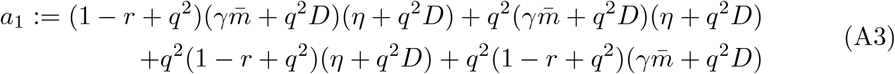

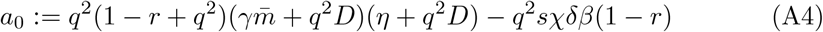

## Appendix B

We set the maximum capacity of our space for cancer cells corresponding to a density of 10^13^ m^3^ in mammalian tissues to *K* = 10^17^ m^*−*2^ ; reference concentrations are set to the following values *c*_0_ = 10^*−*6^ *µ*g*/*m^3^ and *a*_0_ = 10^*−*6^. The values of all the parameters of our model are in Table 1.

**Table B1:** Parameters estimate.

| Description | Dimensional Value | Dimensionless | Reference |
| --- | --- | --- | --- |
| **Motility parameters** |  |  |  |
| $d$ | $10^{-14} \text{ m}^2 \text{ s}^{-1}$ | 1 | Owen et al. Owen and Sherratt (1997) |
| $D$ | $10^{-10} \text{ m}^2 \text{ s}^{-1}$ | $10^4$ | |
| $\chi$ | $10^{-3} \text{ m}^2 \text{ s}^{-1}$ | $10^5$ | Owen et al. Owen and Sherratt (1997) |
| **Degradation/decay rates** |  |  |  |
| $\beta$ | $10^{-6} \text{ m}^3 \text{ g}^{-1} \text{ s}^{-1}$ | $10^{-6}$ | Experimental data |
| $\gamma$ | $10^{-16} \text{ m}^3/\text{s}/$ | $10^7$ | Malier et al Malier et al (2021) |
| $\eta$ | $10^{-4} \text{ s}^{-1}$ | 100 | Knútsdóttir et al. Knútsdóttir et al (2014) |
| **Production/growth rates** |  |  |  |
| $\alpha$ | $10^{-6} \text{ s}^{-1}$ | 1 | Champlain et al. Chaplain and Matzavinos (2005) |
| $\delta$ | $10^{-22} \text{ m}^2 \text{ s}^{-1}$ | $10^5$ | Knútsdóttir et al. Knútsdóttir et al (2014) |
| $s$ | $10^{-2} \text{ g}/\text{m}^3\text{s}$ | $10^{10}$ | Tanaka-Nozaki et al Tanaka-Nozaki et al (2001) |

## Appendix C

In this appendix, we will justify the hypothesis that a population of macrophages metabolizing chemotherapy will lead to a first order decay law. We will consider macrophages as solid volumes *V*_*i*_ present in the domain Ω. The equation governing the evolution of the chemotherapy *c*(**x**, *t*) is the following

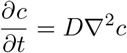

We can integrate this equation on the total volume *V* of the space Ω, which reads

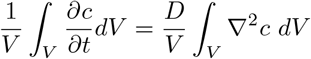

the last part of the equation could be transformed using Ostrogradsky theorem and leads to

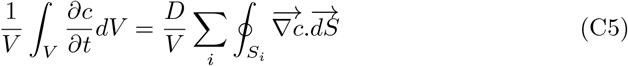

where the Neumann boundary conditions have been used for *∂*Ω. The flux of the chemotherapy at the macrophage surface is at the origin of the *γ* term used in the continuous model of section 2. Here macrophages behave like sinks continuously refilled by diffusion. In spherical coordinates, we can express the equation driving the diffusion of chemotherapy at the surface of one macrophage as follows

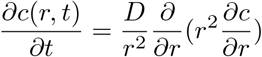

if we consider the size of the macrophage to be *R*_0_, a solution is

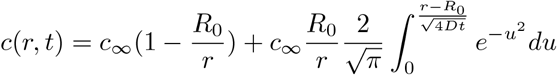

which is valid for *r* ≥ *R*_0_. We can evaluate the flux of chemotherapy at the surface of macrophages with the following formula

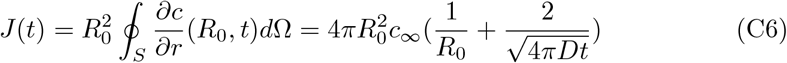

If we consider the mean concentration of 5-FU

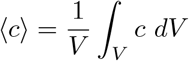

We can insert (C6) in (C5) by considering that density of macrophages is small authorizing to use the equation (C6) by replacing *c* by ⟨*c*⟩. We obtain then for 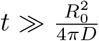 the following first order degradation law

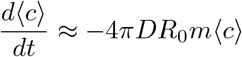

Where 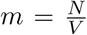 is the density of macrophages. The minus sign coming from the fact that the flux is inward from the macrophages point of view. This equation authorizes to link the *γ* factor of the continuous model and the parameters of the agent-based simulations

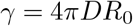

